# Biochemical and structural characterization of a highly active branched-chain amino acid aminotransferase from *Pseudomonas* sp. for efficient biosynthesis of chiral amino acids

**DOI:** 10.1101/559120

**Authors:** Xinxin Zheng, Yinglu Cui, Tao Li, Ruifeng Li, Lu Guo, Defeng Li, Bian Wu

## Abstract

Aminotransferases (ATs) are important biocatalysts for the synthesis of chiral amines because of their capability of introducing amino group into ketones or keto acids as well as their high enantioselectivity, high regioselectivity and no requirement of external addition of cofactor. Among all ATs, branched-chain amino acid aminotransferase (BCAT) can reversibly catalyse branched-chain amino acids (BCAAs), including _L_-valine, _L_-leucine, and _L_-isoleucine, with α-ketoglutaric acid to form the corresponding ketonic acids and _L_-glutamic acid. Alternatively, BCATs have been used for the biosynthesis of unnatural amino acids, such as _L_-tert-leucine. In the present study, the BCAT from *Pseudomonas* sp. (*P*sBCAT) was cloned and expressed in *Escherichia coli* for biochemical and structural analyses. The optimal reaction temperature and pH of *P*sBCAT were 40 °C and 8.5, respectively. *P*sBCAT exhibited a comparatively broader substrate spectrum, and showed remarkably high activity with _L_-leucine, _L_-valine, _L_-isoleucine and _L_-methionine with activities of 105 U/mg, 127 U/mg, 115 U/mg and 98 U/mg, respectively. Additionally, *P*sBCAT had activities with aromatic _L_-amino acids, _L_-histidine, _L_-lysine, and _L_-threonine. To analyse the catalytic mechanism of *P*sBCAT with the broad substrate spectrum, the crystal structure of *P*sBCAT was also determined. Finally, conjugated with the ornithine aminotransferase (OrnAT) from *Bacillus subtilis,* the coupled system was applied to the preparation of _L_-tert-leucine with 83% conversion, which provided an approximately 2.7-fold higher yield than the single BCAT reaction.

**IMPORTANCE:** Despite the enormous potential of BCATs, the vast majority of enzymes still lack suitably broad substrate scope and activity, thus new sources and novel enzymes are currently being investigated. Here, we described a previously uncharacterized *P*sBCAT, which showed a surprisingly wide substrate range and was more active towards BCAAs. This substrate promiscuity is unique for the BCAT family and could prove useful in industrial applications. Based on the determined crystal structure, we found some differences in the organization of the substrate binding cavity, which may influence the substrate specificity of the enzyme. Moreover, we demonstrated efficient biocatalytic asymmetric synthesis of _L_-tert-leucine using a coupling system, which can be used to remove the inhibitory by-product, and to shift the reaction equilibrium towards the product formation. In summary, the structural and functional characteristics of *P*sBCAT were analysed in detail, and this information will play an important role in the synthesis of chiral amino acids and will be conducive to industrial production of enantiopure chiral amines by aminotransferase.

## 1. Introduction

Enantiopure chiral amines are important building blocks in the synthesis of optically active pharmaceuticals, fine chemicals, agrochemicals and bioactive natural products (1–3) due to their structural diversity and functional versatility. The impressive number of applications of chiral amines has stimulated a great deal of innovation in synthetic methodologies for their preparation (4). Within the expanding biocatalytic toolbox for amine synthesis, ATs represent an attractive alternative with excellent optical purity, quantitative yields and high enantioselectivity (5). In general, ATs catalyse the amino group transfer from an amino donor to a ketone or aldehyde amine acceptor with pyridoxal 5′-phosphate (PLP) as cofactor. The first step of the reaction is the transfer of the amino group to the cofactor covalently bound to an active site lysine via a Schiff base, resulting in the formation of pyridoxamine-5’-phosphate (PMP) and release of the amine donor as a ketone or aldehyde. The second step is the regeneration of PLP with the transfer of the amino group to the amine acceptor, which is released as a chiral amino if the acceptor is a ketone or a primary amine if the acceptor is an aldehyde (6).

PLP-dependent enzymes of aminotransferases can be grouped into four subgroups: α family, β family, _D_-alanine transaminase family, and alanine racemase family (7). Among the four subgroups, all ATs belong to the α family except for BCATs and _D_-amino acid aminotransferases (DATs), which belong to the _D_-alanine aminotransferase family (8). Structural similarity and sequence alignment analysis conducted by the Protein Family Database (Pfam) suggest that ATs are divided into six subgroups or classes, with classes I and II including _L_-aspartate aminotransferases (AspATs) and _L_-aromatic amino acid aminotransferases (AroATs), class III ω-transaminases (ω-ATs), class IV BCATs and DATs, class V _L_-serine aminotransferases, and class VI sugar aminotransferases (8, 9).

Among all ATs, BCATs are specific for proteinogenic BCAAs - leucine, isoleucine and valine, to α-ketoglutaric acid to form the respective branched-chain α-ketonic acids and glutamic acid (10). Alternatively, BCATs have been used for the biosynthesis of unnatural amino acids, including _L_-tert-leucine (4). Incorporation of unnatural amino acids into peptide drug serves an important strategy in the synthesis of biologically active compounds with better bioavailability and enhanced potency of the modified target molecules (11). The crucial amino acids in this regard, _L_-tert-leucine, is extensively used for templates in asymmetric synthesis based on its oxazolidine moiety. In addition, _L_-tert-leucine itself is an ingredient of several pharmaceutical developmental products, such as anti-HIV protease inhibitors or tumour-fighting agents (12, 13).

The biosynthesis of _L_-tert-leucine has been achieved by coupling BCAT with several enzymes, including ornithine-δ-aminotransferase (OAT) (14), AspAT/pyruvate decarboxylase (PDC) (15), ω-AT (16), and GluDH/FDH (17). However, the crucial setbacks in the scale-up of the most BCTA coupling reaction include low catalytic efficiency, cumbersome steps and high cost. It is apparent that the pool of enzymes effectively suitable for these biotransformations is limited, and new sources and novel enzymes are currently being investigated.

Here, we successfully expressed and purified BCAT from *Pseudomonas* sp. (*P*sBCAT) and described enzymatic properties, including optimal reaction pH and optimal reaction temperature. The substrate specificity and kinetic parameters of several typical specific substrates were also determined. To further study its structure-function feature, the three-dimensional crystal structure combined with PLP was determined. In addition, compared with *Bs*BCAT, *Cg*BCAT and *Ec*BCAT, *P*sBCAT exhibited highest activities in biosynthesis _L_-tert-leucine, and the feasibility of _L_-tert-leucine synthesis was also verified.

## 2. Materials and methods

### Sequence similarity networks (SSN) analysis

A sequence similarity network of the BCAT family proteins [57,443 sequences, PFam01063 from Pfam release 31.0] (18) was created using EFI-EST web tools (19). In the 1^st^ round, 57,443 sequences from Pf01063 were clustered by 90% identity to 30,927 sequences for alignment, and a network with an E-value cut-off 10^-190^ was created and visualized using the default layout in Cytoscape 3.7.1 (20). To filter out highly conservative sequences and mine sequences with different substrates specificity, nodes with multiple InterPro (21) annotations containing more than 40 sequences from at least 2 prokaryotic genus and not grouped in networks were chosen as input sequences for the next round of analysis. The 2^nd^ network was built with an E-value cut-off 10^-75^ to exclude ungrouped nodes. The results were displayed in Cytoscape using 50% identity RepNodes to obtain higher resolution. The 2^nd^ network grouped the RepNodes into 3 clusters. Sequences in 3 groups were retrieved for the 3^rd^ analysis. The third network had an E-value cut-off 10^-3^ and was coloured by family. Families with less than 3 nodes were deleted.

### Cloning, expression and purification of BCATs and *Bs*OrnAT

The *ilvE* gene of *P*sBCAT (NCBI Accession NO. WP_015095447) was codon-optimized and synthesized with a N-terminal His-tag by General Biosystems. The *P*sBCAT gene was inserted into NdeI and XhoI sites of the pET-28a expression vector. The *ilvE* gene encoding *Bacillus subtilis* BCAT (*Bs*BCAT, Genebank Accession NO. CP017112), *Corynebacterium glutamicum* BCAT (*Cg*BCAT, Genebank Accession NO. CP025533) and *Escherichia coli* BCAT (*Ec*BCAT, Genebank Accession NO. LR134248) were amplified using corresponding primers from the genomes, and the *rocD* gene encoding *Bacillus subtilis* OAT (*Bs*OrnAT, Genebank Accession NO. CP019662) was amplified. The amplified products were assembled into the corresponding position of pET-28a using the Gibson assembly method and then transformed into *E. coli* Top10 for screening. All primers used above are presented in Table 1. Finally, the target plasmids were verified by sequencing (Tsingke, Beijing, China).

**Table 1.**
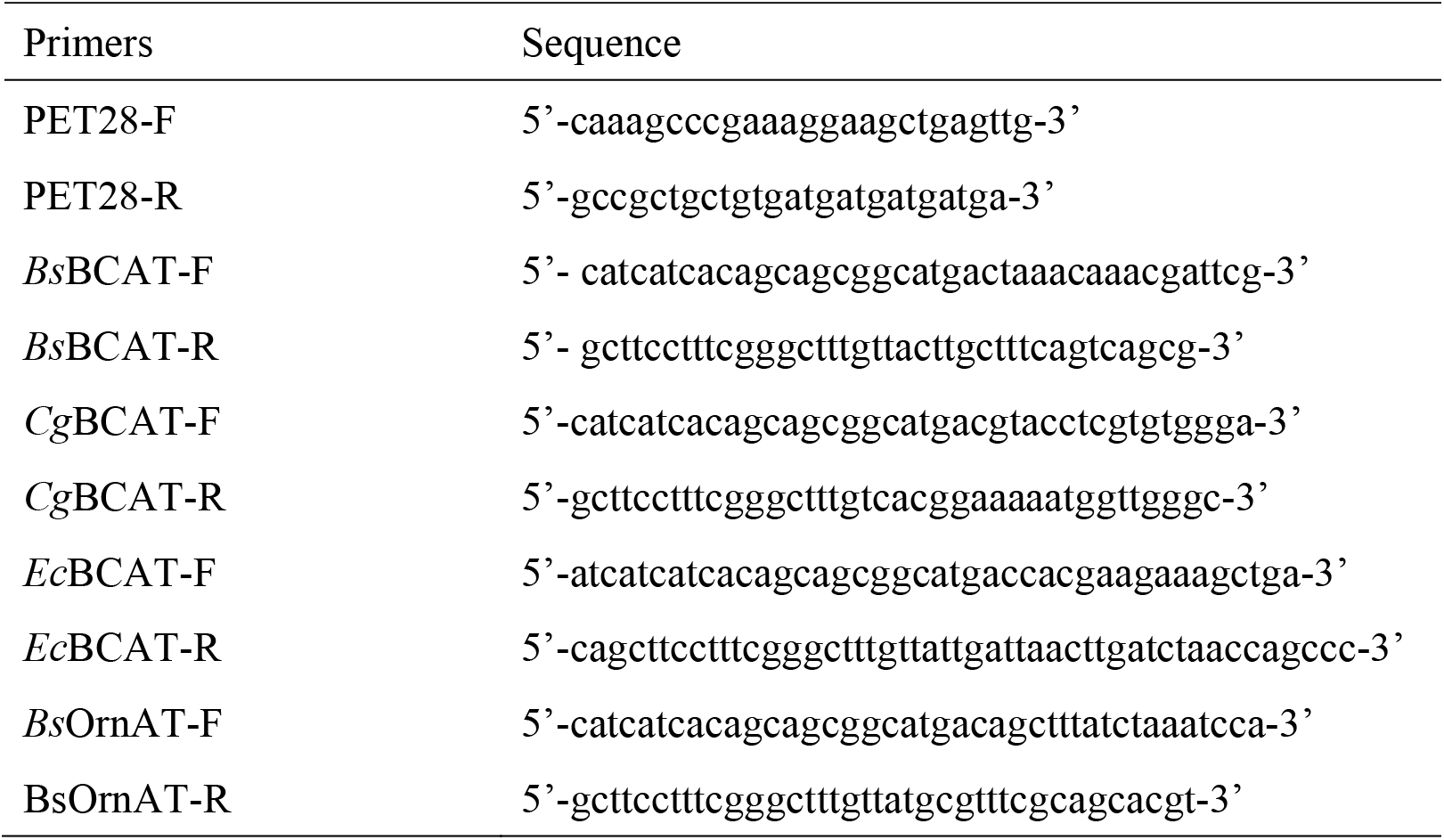
Primers used for gene cloning

*E. coli* BL21(DE3) carrying the recombinant plasmid was cultivated at 30 °C in 1 L auto-induction cultures (10 g/L tryptone, 5 g/L yeast extract, 3.3 g/L ammonium sulfate, 6.8 g/L monopotassium phosphate, 17.9 g/L disodium hydrogen phosphate, 0.55 g/L glucose, 2.1 g/L lactose, 0.49 g/L magnesium sulfate, 2.0 mL/L glycerol) containing 50 *μ*g/mL kanamycin for the expression of recombinant proteins. After 20 h of induction, the cells were harvested, resuspended in 20 mM phosphate buffer (pH=7.5) containing 0.5 M sodium chloride and 20 mM imidazole, and disrupted by sonication. The crude cell extract was centrifuged for 60 min at 12 000 rpm, and supernatants were prepared for further purification. Crude enzyme solution was loaded into Ni-NTA obtained from GE and washed with 50 mL solution buffer (pH=7.5) containing 20 mM imidazole, 20 mM phosphate and 0.5 M sodium chloride. Then, the proteins were eluted with elution buffer (pH=7.5) containing 0.5 M imidazole, 20 mM phosphate, and 0.5 M sodium chloride and desalted with a phosphate buffer (pH=7.5) containing 20 mM phosphate. Finally, the molecular weight of *P*sBCAT was analysed by 12% SDS-PAGE. Purified enzymes were stored in a −20 °C refrigerator for further research. All chemicals, including antibiotics, medium material, and imidazole, were obtained from Sigma-Aldrich.

### Enzyme activity assay

The activity of recombinant *P*sBCAT was determined using _L_-valine as an amino donor and α-ketoglutaric acid as the amino receptor. The reaction was performed at 37 °C and pH 7.5 (20 mM phosphate buffer) with 10 mM _L_-valine, 4 mM α-ketoglutaric acid, 10 μM PLP and an appropriate amount purified enzyme in a total volume of 1 mL. Reactions were terminated by 1 M HCl, and the concentration of α-ketoisovaleric acid was analysed using a C_18_ column with a Shimadzu HPLC system at 210 nm. Separation of the target component was achieved through an isocratic elution with a mixture of 5% acetonitrile and 95% water (0.1% formic acid) at a flow rate of 1.0 mL/min. One unit of enzyme activity was defined as the amount of enzyme that can produce 1 *μ*mol of α-ketoisovaleric acid per min. All experiments were performed in triplicate.

### Effect of pH, temperature and PLP on *P*sBCAT activity

The optimum pH for the reaction with 10 mM _L_-valine, 4 mM α-ketoglutaric acid and 10 *μ*M PLP was determined at 37 °C as described above using the following buffers: 50 mM Na_2_HPO_4_-citric acid buffer (pH 5.0-7.0), 50 mM Tris-HCl buffer (pH 7.0-9.0) and 50 mM Gly-NaOH buffer (pH 9.0-10.5). The optimal temperature was estimated in 50 mM Tris-HCl buffer (pH 8.5) with 10 mM _L_-valine, 4 mM α-ketoglutaric acid and 10 *μ*M PLP under different temperature (25~55 °C). The effect of PLP concentration on activity was tested at 37 °C by performing 10 mM _L_-valine and 4 mM α-ketoglutaric acid reactions in 50 mM Tris-HCl buffer (pH 8.5) (22).

### Substrate specificity

The substrate specificity of the *P*sBCAT for amino acids or amines as amino donors was investigated using α-ketoglutaric acid as the amino acceptor. The reaction mixture consists of 10 mM amino donors, 4 mM α-ketoglutaric acid, and 10 *μ*M PLP in 50 mM Tris-HCl (pH 8.5) at 37 °C. Reactions were terminated by 1 M HCl, and the reduction of α-ketoisovaleric acid was analysed by HPLC.

### Kinetic analysis

For steady-state kinetic analysis, apparent kinetic constants of *Ps*BCAT for _L_-valine, _L_-leucine, _L_-isoleucine and _L_-tert-leucine ranging from 0 to 50 mM were determined at a fixed concentration of α-ketoglutaric acid in 1-mL reaction mixtures containing 50 mM Tris-HCl (pH 8.5) and 10 μM PLP at room temperature. For _L_-glutamic acid, kinetic constants were measured simultaneously but with a fixed concentration of α-ketoisovaleric acid as amino acceptor. In the reverse direction, apparent kinetic constants of α-ketonic acids were measured as well. Apparent *K_m_* and *V_max_* values were calculated by fitting the Michaelis–Menten equation to the experimental data using the non-linear least-squares fit mode of Origin 9.0.

### Crystallization, data collection and structure determination

Crystals of *P*sBCAT were obtained using the hanging-drop vapour-diffusion method at 16°C by mixing equal volumes of protein (12 mg/mL) and reservoir solution that contained 0.25 M ammonium citrate dibasic and 20% (w/v) PEG 3350. The crystals were soaked in reservoir solution supplied with an additional 20% (v/v) glycerol and then flash-frozen in liquid nitrogen. The diffraction datasets were collected at the Shanghai Synchrotron Radiation Facility (SSRF, China) on a beamline BL19U1 with a wavelength of 0.9789 Å at 100 K. The data were processed with XDS (23) and CCP4 (24). Initial phases were obtained by molecular replacement with PHASER (25) using the structure of branched-chain aminotransferase from *Deinococcus radiodurans* (PDB entry 3UYY) (28) as a search model. The model of *P*sBCAT was automatically and manually built by PHENIX.AUTOBUILD (26) and COOT (27), respectively, and refined with PHENIX (26). The data collection and refinement statistics are summarized in Table 2. The coordinates and structure factors for *P*sBCAT have been deposited in the Protein Data Bank under accession 6JIF.

**Table 2.** Statistics of crystal diffraction and structural refinement of *P*sBCAT

| Crystal diffraction statistics | Value |
| --- | --- |
| Wavelength | 0.98 |
| Resolution range | 47.86-1.70 (1.76-1.70) |
| Space group | P 21 21 21 |
| Unit cell | 71.83 95.72 109.82 90 90 90 |
| Total reflections | 1071694 (75160) |
| Unique reflections | 83772(6091) |
| Multiplicity | 12.8 (12.3) |
| Completeness (%) | 99.9 (99.0) |
| Mean I/sigma (I) | 16.57 (2.86) |
| Wilson B-factor | 20.87 |
| R-merge | 0.11 (0.982) |
| R-meas | 0.11 (1.067) |
| R-pim | 0.03134 (0.29) |
| CC1/2 | 0.999 (0.84) |
| CC* | 1 (0.96) |
| Reflections used in refinement | 83754 (8230) |
| Reflections used for R-free | 2000 (197) |
| R-work | 0.19 (0.28) |
| R-free | 0.23 (0.33) |
| CC (work) | 0.96 (0.82) |
| CC (free) | 0.93 (0.71) |
| Number of non-hydrogen atoms | 5927 |
| macromolecules | 5158 |
| ligands | 30 |
| solvent | 739 |
| Protein residues | 667 |
| RMS (bonds) | 0.01 |
| RMS (angles) | 1.05 |
| Ramachandran favored (%) | 97.44 |
| Ramachandran allowed (%) | 2.26 |
| Ramachandran outliers (%) | 0.30 |
| Rotamer outliers (%) | 0.00 |
| Clashscore | 3.42 |
| Average B-factor | 25.53 |
| macromolecules | 24.16 |
| ligands | 19.06 |
| solvent | 35.36 |
| Number of TLS groups | 13 |

Statistics for the highest-resolution shell are shown in parentheses.

### Biosynthesis of _L_-tert-leucine

To investigate the thermodynamic equilibrium of _L_-tert-leucine formation catalysed by *Ps*BCAT, reactions were performed with various concentrations of _L_-glutamic acid (20 mM, 50 mM, 100 mM) in 1 mL of 50 mM Tris-HCl buffer (pH 8.5) containing 20 mM trimethylpyruvic acid and 10 *μ*M PLP and appropriate amount purified enzyme of *P*sBCAT. To shift the unfavourable equilibrium, *Ps*BCAT coupling to *Bs*OrnAT was applied to synthesise _L_-tert-leucine. The reaction mixture contained 0.5 M Tris-HCl buffer (pH 8.5), 0.1 M of trimethylpyruvic acid, 20 mM sodium glutamate, 80 mM _L_-ornithine monohydrochloride, 20 *μ*M PLP and appropriate amounts of purified *P*sBCAT and *Bs*OrnAT enzyme in a total volume of 5 mL at 30 °C. The concentration of _L_-tert-leucine was analysed by HPLC after derivatized using dinitrofluorobenzene (DNFB).

## 3. Results

### Cloning, expression and purification of BCATs

To discover novel BCATs with high activities, BCAT sequences with large sequence diversity were analysed. Three iterative analyses were performed to remove highly conserved BCAT sequences and filter out sequences with low occurrence. After the 3^rd^ round of analysis, a 788-node network was obtained (Figure 1). The representative sequences of the most concentrated clusters (the BCATs from *E. coli*, *Bacillus subtilis* and *Pseudomonas* sp.) were chosen for further analysis. In addition, the well-studied BCAT from *Corynebacterium glutamicum* was also selected. A homology search was conducted based on BCAT from *E. coli* using the blast program against the GenBank database. The BCAT sequences from *Bacillus subtilis*, *Corynebacterium glutamicum* and *Pseudomonas sp.* exhibit 25%, 13% and 30% similarities with that of *E. coli*, representatively.

**Figure 1.**
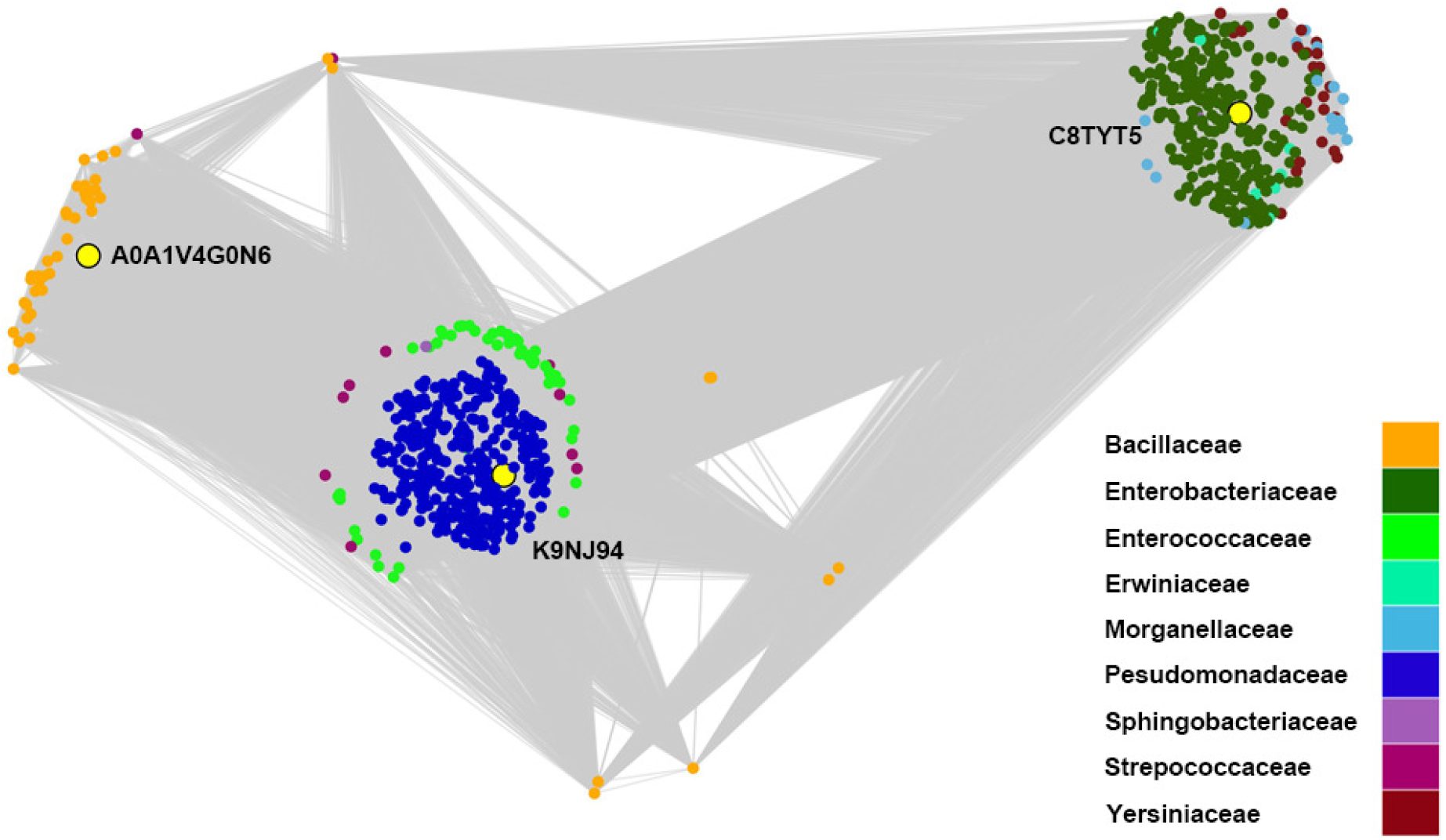
Representation of the BCATs as a sequence similarity network. In this network, nodes represent individual sequences, and edges represent the alignment score from pairwise alignments. Longer edges represent lower scores. Three sequences were selected for further investigation from each cluster. Selected sequences are displayed in yellow with a black border.

The BCAT proteins were produced primarily as soluble protein. The expression of *P*sBCAT was better than other BCATs. *Bs*BCAT, *Ec*BCAT followed, and *Cg*BCAT was the least. The purified BCATs of *Bs*BCAT, *Cg*BCAT, *Ec*BCAT and *P*sBCAT were electrophoretically homogenous as shown on SDS-PAGE with apparent molecular weights of approximately 40.3 kDa, 28.2 kDa, 35.5 kDa, and 37.3 kDa (Figure 2), respectively. Molecular weights of *P*sBCAT and *Ec*BCAT are similar with the *P*sBCAT homologous BCATs, such as *Dr*BCAT (28), *Sm*IlvE (29) and *Mt*IlvE (30) at 38~39 kDa, but higher than that from *Methanococcus aeolicus* (31) with molecular masses of 31.8 kDa for each monomer of the hexamers. The molecular weight of *Bs*BCAT is similar with that of mammalian BCATs, which are commonly found as dimers with molecular masses of 41.3 and 43 kDa for the hBCATm (32) and hBCATc (33) monomers, respectively. Among the four BCATs, *P*sBCAT exhibits highest activity towards _L_-tert-leucine at 7±0.7 U/mg at 37 °C and pH 8.5. The activities of *Bs*BCAT, *Cg*BCAT, and *Ec*BCAT were 1±0.1 U/mg, 2±0.3 U/mg, and 4±0.3 U/mg, respectively. Therefore, *P*sBCAT was chosen for further analysis.

**Figure 2.**
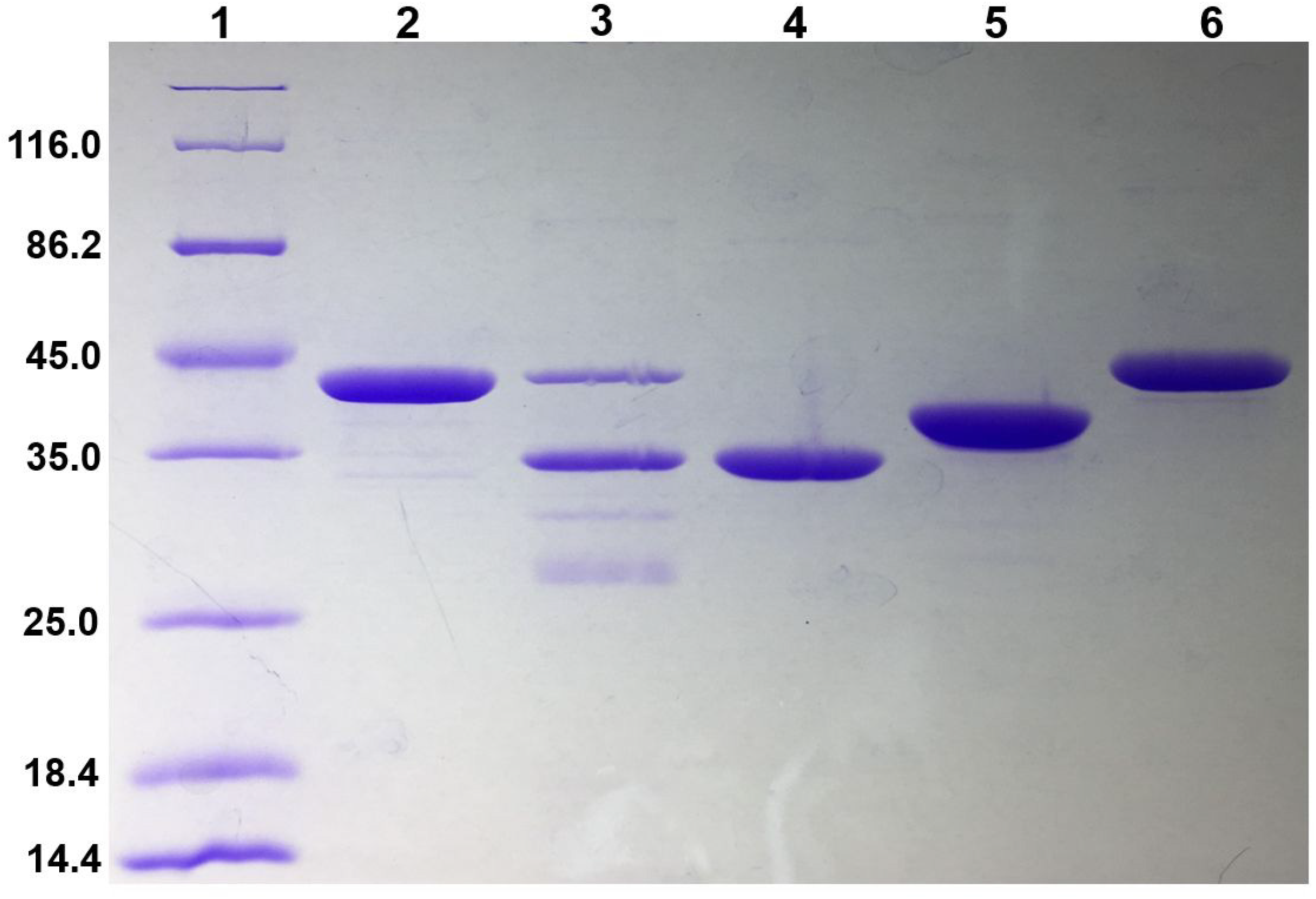
SDS–PAGE of purified BCATs. Lane 1, protein marker; lane 2, 3, 4, 5, and 6 purified *B*sBCAT, *C*gBCAT, *E*cBCAT, *P*sBCAT and *B*sOrnAT, respectively.

### Effect of pH and temperature on *P*sBCAT activity

The effect of pH on *P*sBCAT activity was examined over pH values ranging from 5.0 to 10.5 at 37 °C (Figure 3A). The highest activity of *P*sBCAT for _L_-valine was observed at pH 8.5 in 50 mM Tris-HCl buffer, while it decreased significantly with pH values less than 7.5 or greater than 10.0 with only approximately 10 % activity or even less. The results were consistent with that of most BCATs, including the thermal BCATs from *Vulcanisaeta moutnovskia* (34) and *Thermoproteus uzoniensis* (35, 36), the optimum pH of which were observed at pH 8.5.

**Figure 3.**
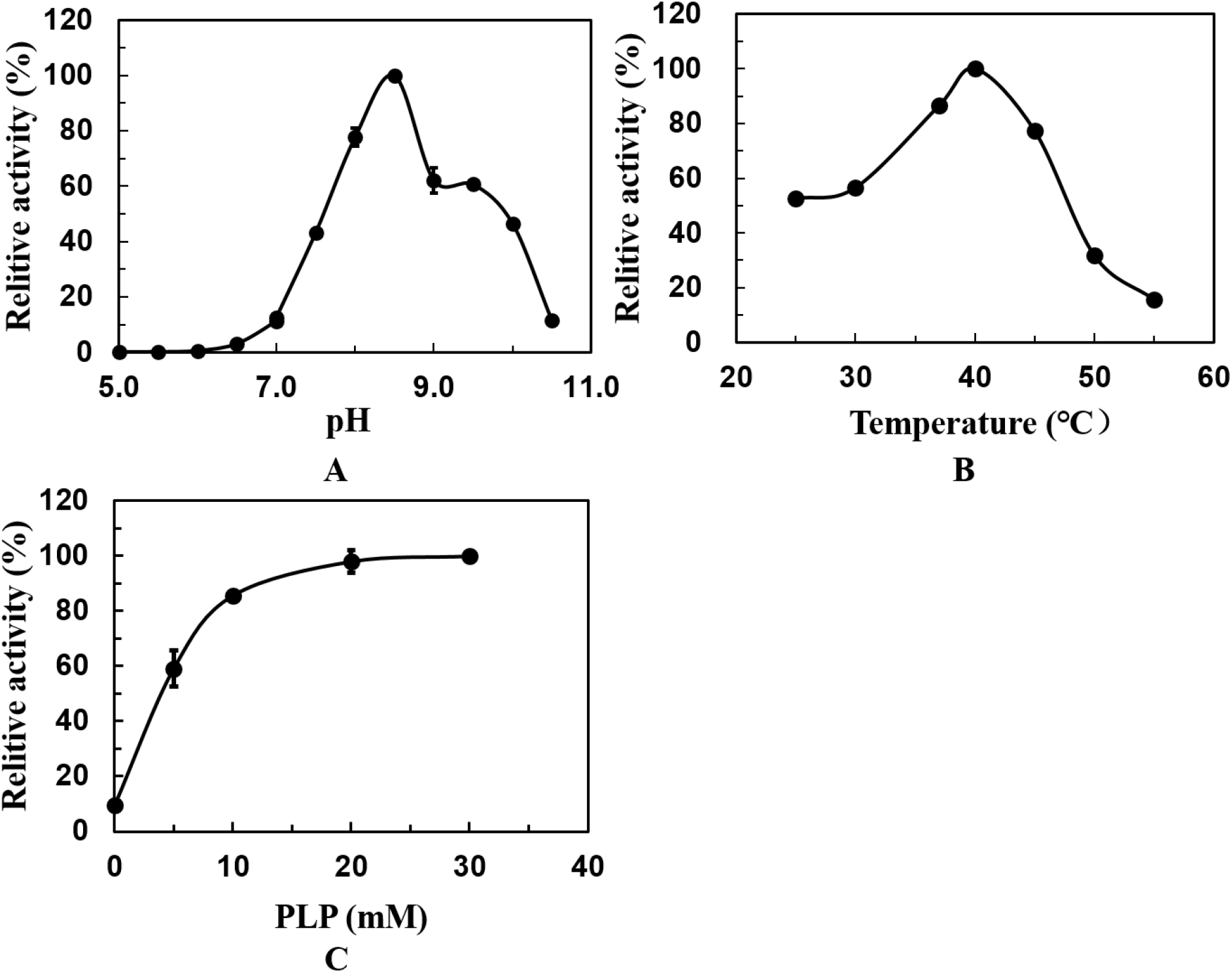
Effects of pH, temperature and PLP on the activities of the *P*sBCAT. (A) The optimal reaction pH of *Ps*BCAT was 8.5. (B) The optimal reaction temperature of *Ps*BCAT was 40 °C. (C) The PLP concentration was 20 at the peak of enzyme activity.

In terms of temperature, *P*sBCAT has an optimal temperature typical of most of BCATs at 40 °C. As shown in Figure 3B, the *P*sBCAT activity decreased very quickly to approximately 30 % at 50 °C, indicating that it was unstable at high temperatures. However, the temperature preference of *P*sBCAT exhibited significant differences compared with thermal BCATs from *Thermoproteus uzoniensis* (35, 36), *Thermococcus* sp. CKU-1 (37) and *Vulcanisaeta moutnovskia* (34), which showed maximum activity at > 90 °C.

### Investigation of PLP requirement of PsBCAT

The purified *P*sBCAT showed low activity when assayed in the absence of PLP but was significantly activated in the presence of PLP (Figure 3C). The effect of PLP concentration on *P*sBCAT activity was investigated using _L_-valine as the substrate, and the Michaelis constant was 3.4 μM. The maximum activity of *P*sBCAT was observed when the PLP concentration reached 20 μM, and no significant activity increase was noted at higher PLP concentrations.

### Substrate specificity and kinetic analysis of *P*sBCAT

To evaluate the substrate specificity of *P*sBCAT, we determined its reactivity toward a series of amine donors (Figure 4). *P*sBCAT exhibited maximum activity toward _L_-leucine followed by _L_-valine, _L_-isoleucine and _L_-methionine with specific activities of 127 U/mg, 115 U/mg, 105 U/mg and 98 U/mg, respectively. Despite the considerably lower activity, *P*sBCAT also showed activities toward _L_-phenylalanine, _L_-alanine and _L_-tert-leucine, but it was inert to _L_-tryptophan, _L_-tyrosine, _L_-threonine, _L_-histidine, _L_-lysine, _L_-theanine and prebalin. Moreover, *P*sBCAT exhibited no activity for amines in propylamine, aniline, and phenylethylamine.

**Figure 4.**
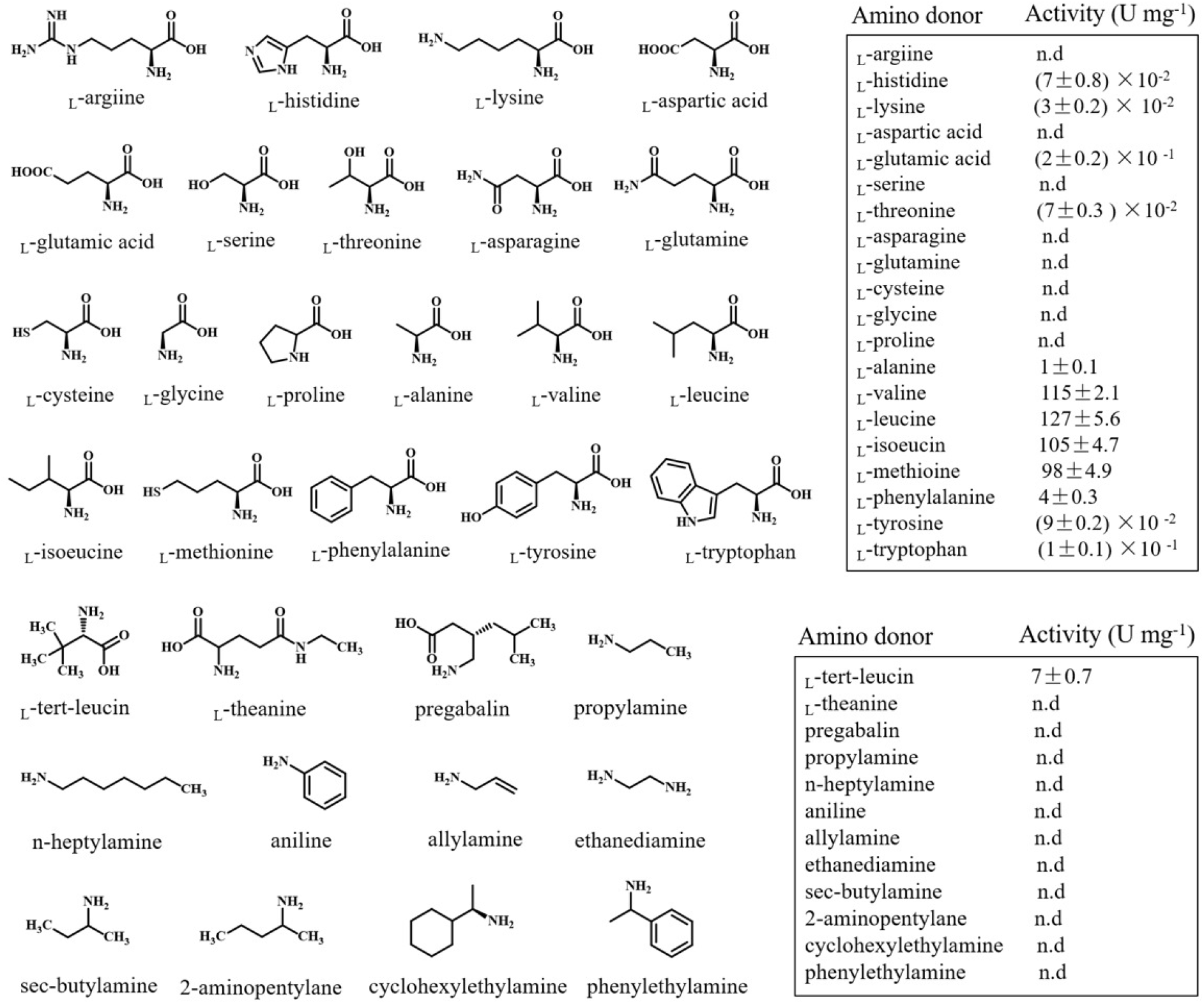
Substrate specificity of *P*sBCAT

The results demonstrate that _L_-leucine, _L_-isoleucine, _L_-valine, _L_-glutamic acid, and _L_-tert-leucine served as suitable substrates. The kinetic parameters were determined for these reactions in two-side directions (Table 3). The effects of the concentration of amino donors on the initial rate followed a typical Michaelis-Menten equation. The *k*_cat_/*K*_m_ value of _L_-leucine was much greater than any others, suggesting that the active site of *Ps*BCAT has evolved to catalyse the transamination reaction for _L_-leucine most effectively. For _L_-tert-leucine transamination, the *K*_m_ value (11.50 mM) was approximately 5-fold higher than that for _L_-leucine, and the *k*cat/*K*m value was much lower (0.90 s^−1^ mM^−1^). Since previous reports have shown that the enzyme activities of aminotransferases were affected by the product or substrate concentrations, product or substrate inhibitions extensively exist among BCATs (10). Kinetics experiments demonstrated that substrate inhibition by _L_-leucine, _L_-isoleucine, _L_-valine, _L_-glutamic acid, and _L_-tert-leucine were not observed up to 50 mM. On the contrary, the ketonic acids exhibit significant inhibition of *Ps*BCAT. For example, the reaction rate increased as the concentration of α-ketoglutaric acid increased up to 5 mM. However, when α-ketoglutaric acid exceeded 5 mM, the reaction rate was decreased. The reaction rate with 10 mM was 55.4% of that with 5 mM α-ketoglutaric acid. The ketonic acid inhibition by α-ketoisovaleric acid was almost the same. The reaction rate with 25 mM was 66.7% of that with 15 mM α-ketoisovaleric acid. As a result, the apparent kinetic parameters of amino acceptors were not obtained.

**Table 3.** Kinetic analysis of *P*sBCAT

| Substrate | $K_m$ (mM) | $K_{cat}$ ($s^{-1}$) | $K_{cat}/K_m$ ($s^{-1} \text{ mM}^{-1}$) |
| --- | --- | --- | --- |
| L-valine | 9.16±1.37 | 220.13±11.97 | 24.41±2.59 |
| L-leucine | 2.27±0.36 | 165.10±10.37 | 74.13±9.24 |
| L- isoleucine | 2.85±0.77 | 175.57±23.95 | 63.91±10.83 |
| L-tert-leucine | 11.50±2.94 | 9.74±0.30 | 0.90±0.22 |
| L- glutamic acid | 18.70±0.64 | 10.53±0.82 | 0.56±0.03 |

### Overall structure of *P*sBCAT

To analyse the catalytic mechanism of *P*sBCAT with a broad substrate spectrum, the crystal structure of *P*sBCAT was determined at a resolution of 1.9 Å (Figure 5A). Structural alignments of *P*sBCAT with type IV class of PLP dependent enzymes, such as *Dr*BCAT (28) (BCAT from *Deinococcus radiodurans,* PDB entry 3UYY), *Sm*IlvE (29) (BCAT from *Streptococcus mutans,* PDB entry 4DQN) and *Mt*IlvE (30) (BCAT from *Mycobacterium tuberculosis,* PDB entry 3HT5) for all Cα atoms are 0.46 Å, 0.50 Å and 0.93 Å, respectively, which indicated that *P*sBCAT shared typical folds of the type IV class of PLP-dependent enzymes.

**Figure 5.**
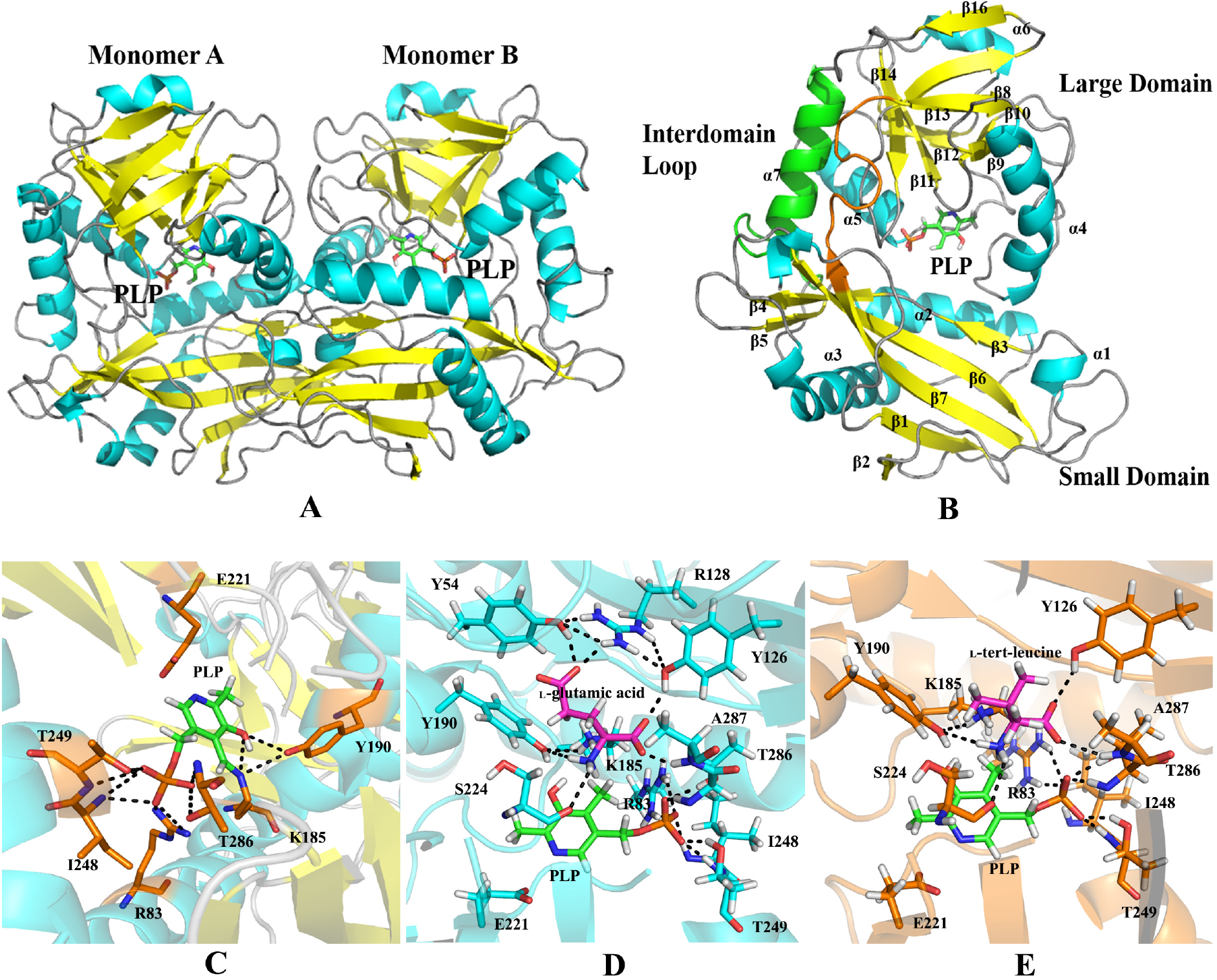
Crystal structures of *P*sBCAT. (A) The *P*sBCAT dimer is shown from an orientation almost perpendicular to the 2-fold axis. The PLP (green stick) is located at the bottom of the active site. (B) The PLP-bound monomer structure of *P*sBCAT is shown with six 7 α-helices (blue) and 16 β-strands (yellow). The monomer structure contains small and large domains linked by the interdomain loop (Val155 to Pro165, shown in orange). The cofactor PLP (green) is located between the two domains. (C) The highly conserved residues combined with PLP in the binding pocket of *P*sBCAT. (D) Interactions among residues, PLP, and the _L_-glutamate molecules. (E) Interactions among residues, PLP, and _L_-tert-leucine molecules.

The asymmetric unit contains one *P*sBCAT homodimer (Figure 5A). Each *P*sBCAT subunit is built up by two separate domains. The first 7 amino acids at the N-terminal region were disordered and invisible in the density map. The small domain (Asn8 to Pro154 and Ile337 to Val340) displays a Rossmann fold and consists of seven β-strands (β1-β7) and three α-helices (α1-α3). Seven β-strands of this domain run in an up-and-down fashion to form a core antiparallel β-sheet with α1-α3 helices on both sides. The large domain (His166 to Pro312) of *P*sBCAT consists of nine β-sheets (β8-β16), which are surrounded by three α-helices (α4-α6). These two domains are linked by α7 helix (residues 313-336) and an interdomain loop (Val155 to Pro165). The interdomain helix is expected to provide a rigid framework for stabilization of the two interacting domains, whereas the interdomain loop is proposed to provide flexibility near the active-site entrance and allow both substrate and product access into and out of the substrate-binding pocket.

### Active site of *Ps*BCAT

The highly conserved residues confer crucial interactions with PLP in the binding pocket of *P*sBCAT, including Arg83, Lys185, Tyr190, Glu221, Ile248, Thr249 and Thr286 (Figure 5C), whereas the corresponding residues are Arg101, Lys204, Tyr209, Glu240, Ile271, Thr272 and Thr314 in *Mt*IlvE (30); Arg100, Lys202, Tyr207, Glu238, Ile266, Thr237 and Thr304 in *Dr*BCAT (28); and Arg81, Lys184, Tyr189, Glu220, Val247, Thr248 and Thr285 in *Sm*IlvE (29). In addition, this positively charged active site pocket is surrounded by residues Phe59, Glu60, Gly61, Tyr126, Arg128, Arg175, Ser224, Ala225, Asn226, Leu245, Gly247 and Gly285.

After docking the ligands into *P*sBCAT, the enzyme-ligand complexes reveal several essential residues for the ligand binding, and some of these residues are conserved in all BCAT enzymes from various species. In the *P*sBCAT-_L_-glutamate complex, Arg128 and Tyr54 confer hydrogen bonds with the γ-carboxylate oxygen atom of _L_-glutamate, whereas Tyr126 and the backbone H atom of Ala287 make hydrogen bonds with the α-carboxylate oxygen atoms. Ser224 and Tyr190 accept hydrogen bonds from the amide hydrogens of _L_-glutamate. The binding mode of _L_-glutamate is similar with that in *Ec*BCAT crystal structure (38). In the *P*sBCAT-_L_-tert-leucine complex, Tyr126 and Ala287 also form hydrogen bonds with the α-carboxylate oxygen atom of _L_-tert-leucine, and the side chain OG atom of Tyr190 and the backbone O atom of Ser224 confer hydrogen-bonding with the amide hydrogens of _L_-glutamate. Remarkably, in both *P*sBCAT-amino acid complexes, Tyr54, Tyr126 and Lys128 make a hydrogen-bonding network to stabilize the carboxylate oxygen atom of the amino acids. The detailed interactions among residues, PLP, and the _L_-glutamate/_L_-tert-leucine molecules are shown in Figure 5D, 5E.

### Application of *P*sBCAT for _L_-tert-leucine biosynthesis

As an important drug intermediate, _L_-tert-leucine is widely used in the synthesis of several pharmaceuticals, such as anti-HIV protease inhibitors or tumour-fighting agents. To measure the chemical reaction equilibrium, syntheses were carried out with various concentrations of _L_-glutamic acid (20 mM, 50 mM, 100 mM) in 1 mL of 50 mM Tris-HCl buffer (pH 8.5) containing 20 mM trimethylpyruvate, 10 *μ*M PLP and appropriate amounts of purified *P*sBCAT. The yields of _L_-tert-leucine were 41%, 52%, and 68%, respectively (Figure 6A, 6B).

**Figure 6.**
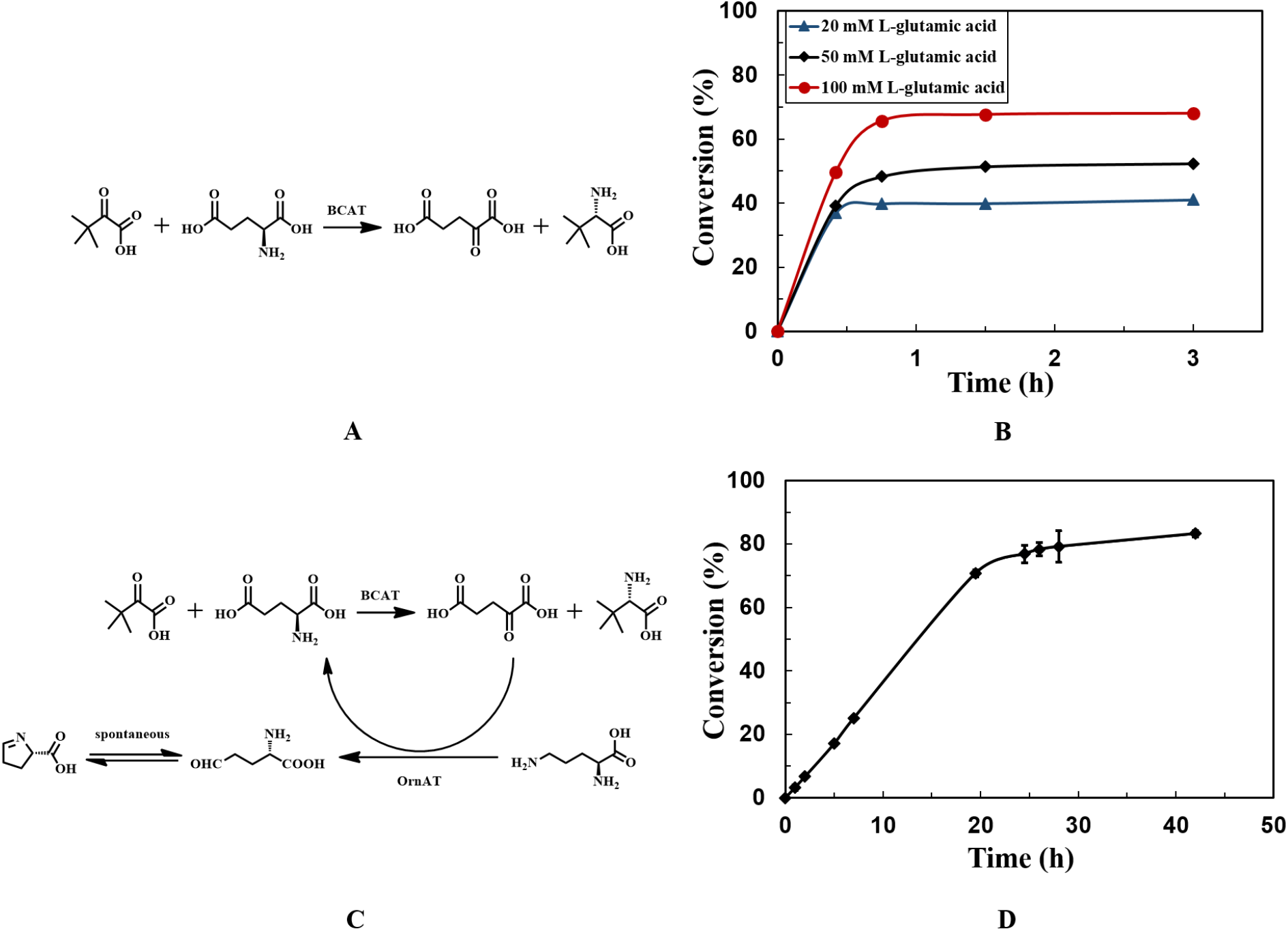
Synthesis of _L_-tert-leucine. (A) The synthesis of _L_-tert-leucine with *P*sBCAT. (B) The reaction was performed in 1 mL of 50 Mm Tris-HCl buffer (pH 8.5) with 20 mM trimethylpyruvate, *P*sBCAT and different concentrations of _L_-glutamic acid (20 mM, 50 mM, and 100 mM). (C) The synthesis of the _L_-tert-leucine with *P*sBCAT coupling BsOrnAT. (D) The reaction was performed in 5 mL of 0.5 M Tris-HCl buffer (pH 8.5) with 100 mM trimethylpyruvate, *P*sBCAT, *Bs*OrnAT; 20 mM _L_-glutamic acid; or 80 mM _L_-ornithine.

Although the concentration of _L_-glutamic acid was up to five times than that of trimethylpyruvate, the substrate cannot be completely transformed. To overcome the product inhibition by α-ketoglutaric acid and shift the unfavourable thermodynamic equilibrium to the synthesis of _L_-tert-leucine, a *Ps*BCAT/*Bs*OrnAT coupling reaction was performed in which _L_-ornithine δ-aminotransferase transferred the δ-amino group from _L_-ornithine to α-ketoglutaric acid, producing _L_-glutamate-γ-semialdehyde and _L_-glutamic acid. The formation of _L_-glutamic acid from α-ketoglutarate was strongly favoured by the spontaneous cyclization of _L_-glutamate-γ-semialdehyde to form Δ^1^-pyrroline-5-carboxylate and drove the reaction to the formation of _L_-tert-leucine (Figure 6C). The synthesis of _L_-tert-leucine were performed with 0.1 M trimethylpyruvate in 5 mL of 0.5 M phosphate buffer (pH 8.5) containing 20 mM _L_-glutamic acid, 80 mM _L_-Ornithine, 20 μM PLP and appropriate amounts of purified *P*sBCAT and BsOrnAT with a yield of 83% at 42 h (Figure 6D).

## Discussion

Aminotransferases are pyridoxal PLP-dependent enzymes that catalyse reversible transamination reactions between amino acids and α-ketoglutaric acid. This enzyme is important for the conversion between inorganic nitrogen and organic nitrogen in living organisms (4).

*P*sBCAT (BCAT from *Pseudomonas* sp.), comprising 340 amino acids, shows many features typical of bacterial BCATs or BCATs from eukaryotes: a comparable molecular weight, similarities in the secondary, tertiary (fold type IV) and quaternary structure, optimal pH value, activity toward BCAAs and _L_-glutamic acid, and inhibition by high concentrations of α-ketonic acids. Meanwhile, *P*sBCAT displays broad but atypical substrate specificity that catalysed the overall transamination reaction of various branched chain amino acids and hydrophobic amino acids when α-ketoglutaric acid was used as an amino accepter. The enzyme has remarkably high activity with _L_-leucine, _L_-valine, _L_-isoleucine and _L_-methionine (127, 115, 105 and 98 U/mg, respectively), whereas other bacterial BCATs only show approximately 10-40 U/mg activities at 37 °C. Additionally, *P*sBCAT shows activities with aromatic _L_-amino acids, _L_-histidine, _L_-lysine, and _L_-threonine, thus displaying broad amino donor specificity.

The molecular mechanism of substrate recognition for BCATs was proposed by Goto based on the analysis of the data on the substrate specificity of *E*cBCAT and structural data (38). The torsional angle of C3-C4-C4’-N of PLP with Lys185 in the *Ps*BCAT crystal structure is 17°, indicating that the internal aldimine bond (Schiff base, C4’=N) is essentially coplanar with the PLP ring to make a conjugated π-system of the PLP-Schiff base. The C4’=N double bond is fixed by the phosphate group, the pyridine hydroxyl group and the pyridine N atom group (Figure 5A). The phosphate group forms hydrogen bonds and salt bridges with Arg83, Ile248, Thr249 and Thr286 and water molecules. The oxygen atom of the PLP pyridine hydroxyl group accepts hydrogen bonds with Tyr190.

BCAT active sites can accommodate two types of substrates with a hydrophobic side chain (branched chain or aromatic amino acids) and with an acidic side chain (acidic amino acids, 2-oxoglutarate) in almost the same positions in the substrate-binding pocket. Phe14, Phe59, Phe159, Tyr190, Thr286, Ala287 are suggested to confer hydrophobic interactions with the hydrophobic or aromatic side chains of the _L_-amino acids, whereas Arg128, Tyr54 and Tyr126 are responsible for the binding of acidic side chain. A comparison of *P*sBCAT and *Dr*BCAT complex structures shows a distinct dynamic loop (Gly156 to Gly162), which is inferred to block the access of the solvent molecules to the active site, thereby preventing the side process of racemization that occurs in the PLP-dependent alanine racemase with the solvent-accessible active site. The Ala157 residue in *P*sBCAT, corresponding to Pro157 in other BCATs, may significantly facilitate the flexibility of the interdomain loop. In addition, Glu197 in BCATs from *Escherichia coli* (38) (PDB ID: 1IYD) and *Thermus thermophilus* (unpublished work) (PDB ID: 1WRV) is suggested to confer electrostatic interactions for substrate binding. However, the truncated side chain of the corresponding residue Ala225 in *P*sBCAT seems to enlarge the large pocket, which might aid in the entry of substrates with bulky moieties. Previous studies have reported _L_-tert-leucine synthesis using BCAT/AspAT/PDC, BCAT/ω-AT, and BCAT/GluDH/FDH coupling systems. In the BCAT/AspAT/PDC coupling system, the formation of _L_-glutamic acid from α-ketoglutaric acid was favoured by the spontaneous decarboxylation of oxaloacetate to form pyruvate (16). Pyruvate decarboxylase (PDC) further performs the non-oxidative thiamine diphosphate-mediated decarboxylation of pyruvate to shift the equilibrium of the reaction (16). The BCAT/ω-AT coupling reaction was also developed to eliminate the product inhibition by simultaneously performing an energetically favourable ω-AT reaction using a two-liquid phase reaction system (17). Recently, for the efficient synthesis of _L_-tert-leucine, BCAT from *Enterobacter* sp. coupling with _L_-glutamate dehydrogenase and formate dehydrogenase was applied. The _L_-glutamate dehydrogenase catalyses the NADPH-dependent amination of α-ketoglutaric acid to yield _L_-glutamic acid, and formate dehydrogenase regenerates NADH using formate (18). Compared with these coupling systems, no requirement of complicated cascade reactions or an expensive external cofactor, such as NADH, renders the BCAT/OAT coupling system promising for industrial applications in the asymmetric synthesis of _L_-tert-leucine. The formation of _L_-glutamic acid from α-ketoglutaric acid was strongly favoured by the spontaneous cyclization of _L_-glutamate-γ-semialdehyde to form Δ^1^-pyrroline-5-carboxylate and drove the reaction to the formation of _L_-tert-leucine. Indeed, *P*sBCAT/OAT coupling reaction produced 83 mM _L_-tert-leucine from 100 mM trimethylpyruvate for 42 h, which provided an approximately 2.7-fold higher yield than the single BCAT reaction.

In conclusion, we determined the crystal structure of branched-chain amino acid aminotransferase from *Pseudomonas* sp. The enzyme is active toward typical BCAT substrates, including _L_-leucine, _L_-isoleucine and _L_-valine, and also shows significant activity with aromatic _L_-amino acids, _L_-histidine, _L_-lysine, and _L_-threonine. The structure of *P*sBCAT is similar to the known structures of bacterial BCATs. However, we found some differences in the organization of the substrate binding cavity, which may influence the substrate specificity of the enzyme. Taking into account a broad range of accepted substrates and detected ability to catalyse asymmetric synthesis of _L_-tert-leucine, this enzyme hold promises in biotechnological applications, and the structural fingerprint will aid in further engineering to increase the activity with amines.

## SUPPLEMENTAL MATERIAL

Supplementary material for this manuscript has been submitted.

## Acknowledgment

This work is supported by the National Natural Science Foundation of China (Grant Nos. 21603013, 31601412, 31822002), the 100 Talent Program grant and Biological Resources Service Network Initiative (ZSYS-012) from the Chinese Academy of Sciences.

## References

1. Ramesh NP. 2013. Biocatalytic synthesis of chiral alcohols and amino acids for development of pharmaceuticals. Biomolecules 3:741–777. https://doi.org/10.3390/biom3040741.

2. Gordon EM, Ondetti MA, Pluscec J, Cimarusti CM, Bonner DP, Sykes, RB. 1982. O-Sulfated .beta.-lactam hydroxamic acids (monosulfactams). Novel monocyclic .beta.-lactam antibiotics of synthetic origin. J Am Chem Soc 104:6053–6060. https://doi.org/10.1021/ja00386a035.

3. Gustafsson D, Elg M, Lenfors S, Börjesson I, Teger-Nilsson AC. 1996. Effects of inogatran, a new low-molecular-weight thrombin inhibitor, in rat models of venous and arterial thrombosis, thrombolysis and bleeding time. Blood Coagul 7:69–79. https://doi.org/10.1097/00001721-199601000-00009.

4. Xue YP, Cao CH, Zheng YG. 2018. Enzymatic asymmetric synthesis of chiral amino acids. Chem Soc Rev 47:1516–1561. https://doi.org/10.1039.C7CS00253J.

5. Hwang BY, Cho B K, Yun H, Koteshwar K, Kim B G. 2005. Revisit of aminotransferase in the genomic era and its application to biocatalysis. J Mol Catal B-Enzym 37:47–55. https://doi.org/10.1016/j.molcatb.

6. Taylor PP, Pantaleone DP, Senkpeil RF, Fotheringham IG. 1998. Novel biosynthetic approaches to the production of unnatural amino acids using transaminases. Trends Biotechnol 16:412–418. https://doi.org/10.1016/S0167-7799(98)01240-2.

7. Jansonius JN. 1998. Structure, evolution and action of vitamin b6-dependent enzymes. Curr Opin Struct Bio 8:759–769. https://doi.org/10.1016/S0959-440X(98)80096-1.

8. Slabu I, Galman JL, Lloyd RC, Turner NJ. 2017. Discovery, engineering and synthetic application of transaminase biocatalysts. ACS Catal 7:8263–8284. https://doi.org/10.1021/acscatal.7b02686.

9. Finn RD, Mistry J, Tate J, Coggill P, Heger A, Pollington JE, Gavin OL, Gunasekaran P, Ceric G, Forslund K, Holm P, Sonnhammer EL, Eddy SR, Bateman A. 2010. The Pfam protein families database. Nucleic Acids Res 38: D211–D222. https://doi.org/10.1093/nar/gkp985.

10. Bezsudnova EY1, Boyko KM, Popov VO. 2017. Properties of bacterial and archaeal branched-chain amino acid aminotransferases. Biochemistry 82:1572–1591. https://doi.org/10.1134/S0006297917130028.

11. Rudat J, Brucher BR, Syldatk C. 2012. Transaminases for the synthesis of enantiopure beta-amino acids. AMB Express. 2:1–10. https://doi.org/10.1186/2191-0855-2-11.

12. Allenmark S, Lamm B. 2001. A useful route to (r)-and (s)-tert-leucine. Chirality 13:43–47. https://doi.org/10.1002/1520-636X(2001)13:1<43::AID-CHIR9>3.0.CO;2-G.

13. Hong EY, Cha M, Yun H, Kim BG. 2010. Asymmetric synthesis of l-tert-leucine and 1-3-hydroxyadamantylglycine using branched chain aminotransferase. J Mol Catal B-Enzym 66:228–233. https://doi.org/10.1016/j.molcatb.2010.05.014.

14. Lehr P, Billich A, Charpiot B, Ettmayer P, Scholz D, Rosenwirth B, Gstach H. 1996. Inhibitors of human immunodeficiency virus type 1 protease containing 2-aminobenzyl-substituted 4-amino-3-hydroxy-5-phenylpentanoic acid: synthesis, activity, and oral bioavailability. J Med Chem 39:2060–2067. https://doi.org/10.1021/jm9508696.

15. Li T, Kootstra AB, Fotheringham IG. 2002. Nonproteinogenic α-amino acid preparation using equilibrium shifted transamination. Org Process Res Dev 6:533–538. https://doi.org/10.1021/op025518x.

16. Seo YM, Yun H. 2011. Enzymatic synthesis of l-tert-leucine with branched chain aminotransferase. J Microbiol Biotechnol 21:1049–1052. https://doi.org/10.1021/op025518x.

17. Park ES, Shin JS. 2015. Biocatalytic cascade reactions for asymmetric synthesis of aliphatic amino acids in a biphasic reaction system. J Mol Catal B-Enzym 121:9–14. https://doi.org/10.1016/j.molcatb.2015.07.011.

18. Gebali ES, Mistry J, Bateman A, Eddy SR, Luciani A, Potter SC, Qureshi M, Richardson LJ, Salazar GA, Smart A, Sonnhammer ELL, Hirsh L, Paladin L, Piovesan D, Tosatto SCE, Finn RD. The Pfam protein families database in 2019. Nucleic Acids Res D427–D432. https://doi.org/10.1093/nar/gky995

19. Gerlt JA, Bouvier JT, Davidson DB, Imker HJ, Sadkhin B, Slater DR, Whalen KL. 2015. Enzyme function initiative-enzyme similarity tool (efi-est): a web tool for generating protein sequence similarity networks. Biochim Biophys Acta 1854:1019–1037. https://doi.org/10.1016/j.bbapap.2015.04.015.

20. Shannon P, Markiel A, Ozier O, Baliga NS, Wang JT, Ramage D, Amin N, Schwikowski B, Ideker T. 2003 Cytoscape: A Software Environment for Integrated Models of Biomolecular Interaction Networks. Genome Res 13:2498–2504. https://doi.org/10.3372/wi.43.43119.

21. Mitchell AL, Attwood TK, Babbitt PC, Blum M, Bork P, Bridge A, Brown SD, Chang HY, Gebali SE, Fraser MI, et al. InterPro in 2019: improving coverage, classification and access to protein sequence annotations. Nucleic Acids Res 47:D351–D360. https://doi.org/10.1093/nar/gky1100.

22. Zheng RC, Tang XL, Suo H, Feng LL, Liu X, Yang J, Zheng YG. 2018. Biochemical characterization of a novel tyrosine phenol-lyase from Fusobacterium nucleatum for highly efficient biosynthesis of l-DOPA. Enzyme Microb Technol 112:88–93. https://doi.org/10.1016/j.enzmictec.2017.11.004.

23. Kabsch W. 2010. XDS. Acta Crystallogr D Biol Crystallogr 66:125–132. pmid: 20124692. https://doi.org/10.1107/S0907444909047337.

24. Winn MD, Ballard CC, Cowtan KD, Dodson EJ, Emsley P, Evans PR, Keegan RM, Krissinel EB, Leslie AG, McCoy A, McNicholas SJ, Murshudov GN, Pannu NS, Potterton EA, Powell HR, Read RJ, Vagin A, Wilson KS. 2011. Overview of the CCP4 suite and current developments. Acta Crystallogr D Biol Crystallogr 67:235–242. pmid: 21460441. https://doi.org/10.1107/S0907444910045749.

25. McCoy AJ, Grosse-Kunstleve RW, Adams PD, Winn MD, Storoni LC, Read RJ. 2007. Phaser crystallographic software. J Appl Crystallogr 40:658–674. https://doi.org/10.1107/S0021889807021206.

26. Adams PD, Afonine PV, Bunkóczi G, Chen VB, Davis IW, Echols N, Headd JJ, Hung LW, Kapral GJ, Grosse-Kunstleve RW, McCoy AJ, Moriarty NW, Oeffner R, Read RJ, Richardson DC, Richardson, Jane SR, Thomas CT, Peter HZ. 2010. PHENIX: a comprehensive Python-based system for macromolecular structure solution. Acta Crystallogr. D Biol. Crystallogr 66:213–221. https://doi.org/10.1107/S0907444909052925.

27. Emsley P, Cowtan K. 2004. Coot: model-building tools for molecular graphics. Acta Crystallogr. D Biol. Crystallogr 60:2126–2132. https://doi.org/10.1107/S0907444904019158.

28. Chen CD, Lin CH, Chuankhayan P, Huang YC, Hsieh YC, Huang TF, Guan HH, Liu MY, Chang WC, Chen CJ. 2012. Crystal Structures of Branched-Chain Aminotransferase from Deinococcus radiodurans Complexes with α-Ketoisocaproate and L-Glutamate Suggest Its Radiation-Resistance for Catalysis. J Bacterio 194:6206–6216. https://doi.org/10.1128/JB.01659-12.

29. Ruan J, Hu J, Yin A, Wu W, Cong X, Feng X, Li S. 2012. Structure of the branched-chain aminotransferase from *Streptococcus mutans*. Acta Crystallogr D Biol Crystallogr 68:996–1002. https://doi.org/10.1107/S0907444912018446.

30. Tremblay LW, Blanchard JS. 2009. The 1.9 Å structure of the branched-chain amino-acid transaminase (IlvE) from *Mycobacterium tuberculosis*. Acta Crystallogr Sect F Struct Biol Cryst Commun 65:1071–1077. https://doi.org/10.1107/S1744309109036690.

31. Xing RY, Whitman WB. 1992. Characterization of amino acid aminotransferases of *methanococcus aeolicus*. J Bacteriol 174:541–548. https://doi.org/10.1128/jb.174.2.541-548.1992

32. Yennawar N, Dunbar J, Conway M, Hutson S, Farber G. 2001. The structure of human mitochondrial branched-chain aminotransferase. Acta Crystallogr D Biol Crystallogr 57:506–515. https://doi.org/10.1107/S0907444901001925.

33. Goto M, Miyahara I, Hirotsu K, Conway M, Yennawar N, Islam MM, Hutson SM. 2005. Structural determinants for branched-chain aminotransferase isozyme-specific inhibition by the anticonvulsant drug gabapentin. J Biol Chem 280:37246–37256. https://doi.org/10.1074/jbc.M506486200.

34. Stekhanova TN, Rakitin AL, Mardanov AV, Bezsudnova EY, Popov VO. 2017. A novel highly thermostable branched-chain amino acid aminotransferase from the crenarchaeon *Vulcanisaeta moutnovskia*. Enzyme and microbial technology 96:127–134. https://doi.org/10.1016/j.enzmictec.2016.10.002.

35. Bezsudnova EY, Stekhanova TN, Suplatov DA, Mardanov AV, Ravin NV, Popov VO. 2016. Experimental and computational studies on the unusual substrate specificity of branched-chain amino acid aminotransferase from *Thermoproteus uzoniensis*. Arch Biochem Biophys 607:27–36. https://doi.org/10.1016/j.abb.2016.08.009.

36. Boyko KM, Stekhanova TN, Nikolaeva AY, Mardanov AV, Rakitin AL, Ravin NV, Bezsudnova EY, Popov VO. 2016. First structure of archaeal branched-chain amino acid aminotransferase from *Thermoproteus uzoniensis* specific for L-amino acids and R-amines. Extremophiles 20:215–225. https://doi.org/10.1007/s00792-016-0816-z.

37. Uchida Y, Hayashi H, Washio T, Yamasaki R, Kato S, Oikawa T. 2014. Cloning and characterization of a novel fold-type I branched-chain amino acid aminotransferase from the hyperthermophilic archaeon *Thermococcus* sp. CKU-1. Extremophiles 18:589–602. https://doi.org/10.1007/s00792-014-0642-0

38. Goto M, Miyahara I, Hayashi H, Kagamiyama H, Hirotsu K. 2003. Crystal structures of branched-chain amino acid aminotransferase complexed with glutamate and glutarate: true reaction intermediate and double substrate recognition of the enzyme. Biochemistry 42:3725–3733. https://doi.org/10.1021/bi026722f.

